# Transcranial direct current stimulation over left inferior frontal cortex improves speech fluency in adults who stutter

**DOI:** 10.1101/259796

**Authors:** Jennifer Chesters, Riikka Möttönen, Kate E. Watkins

## Abstract

Stuttering is a neurodevelopmental disorder affecting 5% of children, and persisting in 1% of adults. Promoting lasting fluency improvement in adults who stutter is a particular challenge. Novel interventions to improve outcomes are required, therefore. Previous work in patients with acquired motor and language disorders reported enhanced benefits of behavioural therapies when paired with transcranial direct current stimulation (tDCS). Here, we report the results of the first trial investigating whether tDCS can improve speech fluency in adults who stutter. Thirty adult men who stutter completed a randomized, double-blind, controlled trial of anodal tDCS over left inferior frontal cortex. Fifteen men received 20 minutes of 1-mA tDCS on five consecutive days while speech fluency was temporarily induced using choral and metronome-timed speech. The other 15 men received the same speech fluency intervention with sham stimulation. We predicted that applying anodal tDCS to the left inferior frontal cortex during speech production with temporary fluency inducers would result in longer-lasting fluency improvements. Speech fluency during reading and conversation was assessed at baseline, before and after the stimulation on each day of the five-day intervention, and at 1 and 6 weeks after the end of the intervention. TDCS combined with speech fluency training significantly reduced the percentage of disfluent speech measured 1 week after the intervention compared with fluency intervention alone. At 6 weeks after the intervention, this improvement was maintained during reading but not during conversation. Outcome scores at both post-intervention time points on a clinical assessment tool (the Stuttering Severity Instrument – version 4) also showed significant improvement in the group receiving tDCS compared with the sham group, in whom fluency was unchanged from baseline. We conclude that tDCS combined with behavioural fluency intervention has the capacity to improve fluency in adults who stutter. tDCS thereby offers a potentially useful adjunct to future speech therapy interventions for this population, for whom therapy outcomes are currently limited.

## Introduction

Developmental stuttering is a neurodevelopmental disorder disrupting the smooth flow of speech, resulting in characteristic speech disfluencies. Developmental stuttering represents a considerable burden to the 1% of adults living with the condition, and to society; it is associated with reduced educational and employment opportunities (Klein *et al*., 2004; O’Brian *et al*., 2011), social anxiety (Iverach *et al*., 2009), and compromised quality of life (Craig *et al*., 2009). Fluency therapies may use techniques for altering speech patterns to reduce overt stuttering (Boberg *et al*., 1994; O’Brian *et al*., 2003). However, fluency improvements do not persist without continued practice, and can be difficult to fully integrate into everyday speech. Furthermore, learning these new speech patterns can affect speech naturalness (Metz *et al*., 1990; O’Brian et al., 2003; Tasko *et al*., 2007), which can reduce the acceptability of these approaches. There is a value, therefore, in developing novel interventions to improve therapy outcomes for adults who stutter.

Transcranial direct current stimulation (tDCS), a non-invasive brain stimulation method, may have potential to improve the outcomes of fluency interventions in people who stutter (Chesters *et al*., 2017). TDCS involves application of a weak electrical current across the head via electrodes placed on the scalp, modulating the resting membrane potential of neurons in the underlying cortex. Anodal tDCS applied over motor cortex enhances cortical excitability (Nitsche *et al*., 2000) and can improve motor learning (Nitsche *et al*., 2003; Stagg *et al*., 2011). When paired with a task, tDCS can increase neural plasticity, and its effects build and stabilize when applied in consecutive daily sessions (Baker *et al*., 2010; Reis *et al*., 2009). Increasingly, tDCS is being investigated as an adjunctive treatment for acquired disorders of motor, language and cognitive functions (Allman *et al*., 2016; Baker *et al*., 2010; Khedr *et al*., 2013; Marangolo *et al*., 2011; Mortensen *et al*., 2016). For example, in a study treating upper limb motor function in stroke patients, tDCS was found to prolong the effects of 9-days of motor training for at least three months (Allman *et al*., 2016). In post-stroke aphasic patients, five days of anodal tDCS over left inferior frontal cortex enhanced naming accuracy, which remained improved for at least one week post intervention (Baker *et al*., 2010). Here, we aimed to evaluate whether lasting fluency improvements could be obtained in a group of adults who stutter by combining tDCS with a 5-day behavioural fluency intervention.

People who stutter can experience near, or complete, fluency by changing the way speech is produced, for example by speaking with a different accent or in time with an external stimulus, such as a metronome or another speaker (so called ‘choral speech’). Altering the auditory feedback associated with speech production can also be effective; for example, feedback that is noisy, or altered in pitch or time (delayed) can result in almost complete fluency in some people (as portrayed in the film *The King’s Speech).* It is important to note, however, that these forms of fluency induction, while very successful at inducing almost complete fluency, are temporary and that disfluency returns typically once the inducer is removed. Although these fluency inducers are of little efficacy therapeutically, for our purposes their effectiveness in achieving immediate and close to complete fluency, with little impact on naturalness, was an important factor. We hypothesized that by applying tDCS while fluent speech was induced in people who stutter, we could facilitate the brain circuits supporting this fluent speech, promoting neuro-plastic changes and thereby produce lasting fluency improvements.

The effectiveness of the temporary fluency inducers described above is consistent with theories that dysfluency in people who stutter is caused by a problem in generating internal timing cues for motor control or sensorimotor integration or both (Alm, 2004; Max *et al*., 2004; Watkins *et al*., 2015). Brain imaging studies of adults who stutter confirm both structural and functional abnormalities in sensorimotor circuits involved in speech production. Specifically, there is consistent evidence of a white matter structural deficits underlying ventral sensorimotor cortex in the left hemisphere (Sommer *et al*., 2002; Watkins *et al*., 2008; see Neef *et al*., 2015, for review). Functionally, there are differences in activation patterns evoked by speech production in people who stutter that reflect both trait and state differences (see recent meta-analyses by Belyk *et al*., 2015, and Budde *et al*., 2014). Of relevance to our study, it is worth noting that the left inferior frontal cortex is functionally abnormal in people who stutter (Fox *et al*., 1996; Kell *et al*., 2009; Neumann *et al*., 2005; Toyomura *et al*., 2011; Watkins *et al*., 2008; Wu *et al*., 1995). In the current study, we placed the anodal electrode over the left inferior frontal cortex covering also the ventral sensorimotor and premotor cortex.

We recruited 30 adult men who stutter to a randomized double-blind controlled trial using tDCS in combination with a behavioural fluency intervention. The behavioural intervention involved temporarily inducing fluency using both choral speech and metronome-timed speech during overt reading, narrative and and conversational speech tasks. We delivered 1-mA of anodal tDCS over the left inferior frontal cortex (IFC) for 20 minutes per day in five consecutive daily sessions. Fluency was assessed 1 and 6 weeks after the five-day intervention. We predicted that fluency intervention when combined with anodal tDCS would result in reduced disfluency, relative to the same fluency intervention with sham stimulation.

## Methods

### Study design and participants

The study had a double-blind, sham-controlled, parallel-group design. A UK community sample of men aged 18–50 years, with at least a moderate stutter (assessed using the Stuttering Severity Instrument – version 4; SSI-4, Riley, 2009), and with English as a first language, were recruited to participate. Exclusion criteria included any disorder of speech, language or communication other than developmental stuttering, sensory impairment, neurological or psychiatric illness, and any safety contra-indication for tDCS. The University of Oxford Central University Research Ethics Committee (MSD-IDREC-C2-2014–013) approved the study. Participants gave informed written consent to participate in the study, in accordance with the Declaration of Helsinki, and with the procedure approved by the committee. The trial was registered on ClinicalTrials.gov (NCT02288598).

### Randomisation and masking

A researcher who was not involved in any aspect of the trial performed the randomisation of participants into the sham and tDCS study arms using blocked randomization (Roberts *et al*., 1998). A block size of four was chosen generating six possible sequences, which were allocated at random. Allocation concealment was achieved by assigning a unique 5-digit code per participant. The code was used to deliver tDCS or sham stimulation using the study mode on the stimulator (http://www.neurocaregroup.com/dc_stimulator_plus.html). Codes remained in sealed sequentially numbered opaque envelopes until allocation. The participants and the researcher who delivered the intervention, assessed the outcomes, and analyzed the data, were masked to group assignment.

### Procedures

### tDCS

In the tDCS study arm, participants received 20 minutes of stimulation at 1 mA, applied during the fluency intervention. The anode was placed over left IFC (centred on FC5 according to the 10–10 EEG electrode placement system), and the cathode over the right supra-orbital ridge (see figure 1A). A Neuroconn direct-current stimulator in study mode was used to deliver tDCS. The electrodes measured 5 cm x 7 cm, and were placed within saline soaked sponges; the anode was placed in portrait orientation, and the cathode in landscape orientation. The same electrode placement was used in the sham stimulation study arm, during which the current was ramped up over 15 seconds, maintained for 15 seconds at 1 mA and ramped down over 15 seconds at the start of the session, followed by brief (3ms) pulses every 55 seconds for 20 minutes. These sham stimulation parameters delivered current at an ineffective dosage. However, the initial ramping of current and the intermittent current pulse ensured effective blinding of participant and researcher.

**Figure 1:**
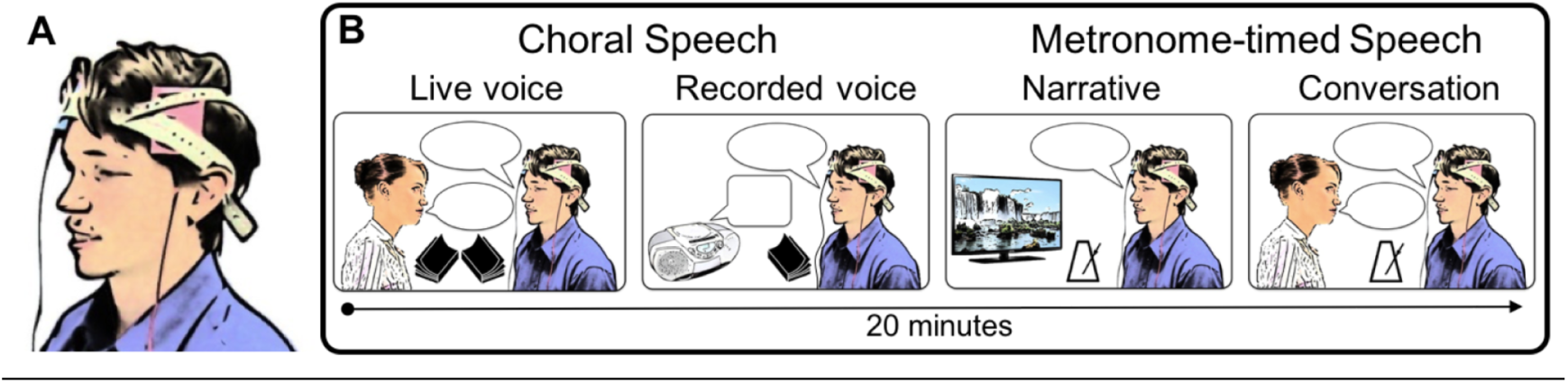
tDCS montage and behavioural tasks used in intervention. A shows the electrode placement montage used to apply tDCS. Anode (red) was placed over left IFC, centred on position FC5 of the 10–10 EEG electrode placement system. Cathode (blue) was placed over the right supra-orbital ridge. B shows the choral speech (live voice and recorded voice) and metronome-timed speech (video narrative and conversation) tasks used in each daily intervention session. 1-mA tDCS was applied concurrently with these tasks for 20 mins in 15 men who stutter. Another 15 men received sham stimulation for the same period. Both researcher and participant were blind to the stimulation condition.

#### Fluency intervention

A registered Speech and Language Therapist delivered the fluency intervention in 20-minute sessions on 5 consecutive days. We used behavioural techniques that induce temporary fluency. We chose these techniques for a maximal and immediate fluency induction because we wanted to be sure that application of tDCS would promote only the fluent state of speech and not the disfluent one. The behavioural techniques were choral speech, and metronome-timed speech (Kiefte *et al*., 2008; Trajkovski *et al*., 2009). Figure 1B illustrates the tasks used in each intervention session. Choral speech involved reading passages at a normal rate in unison first with a live voice and second with an audio-book recording. Metronome-timed speech involved speaking in time with an external audio metronome to produce spontaneous narratives of silent cartoon films followed by conversation with the researcher on randomly selected topics (e.g. a recent holiday). The metronome rate increased from 140 to 190 beats per minute across the five days. Participants were instructed to indicate if the rate exceeded a comfortable speaking rate at any time. In this case, the metronome was slowed to a comfortable rate, and maintained at this rate for the remaining days. Speech disfluency during reading and conversation was measured at baseline, before and after the intervention on each day, and at 1 week and 6 weeks post-intervention.

### Outcome measures

The primary outcome measure for the trial was change from baseline proportion of stuttering in speech samples taken at 1 and 6 weeks post intervention. A baseline percentage of disfluent syllables ( %ds) was estimated for two speech samples taken during reading and conversation on two separate days and averaged to give a stable estimate. The same measurement was taken post-intervention and the change from baseline at each time point calculated by subtracting the baseline from the post-intervention estimates. The primary outcome measure was overall change in fluency; we also analysed data from each task separately to explore whether the speaking situation produced different effects. We defined disfluent syllables as those containing repetition or prolongation of a speech sound, or where a tense pause or ‘block’ occurred prior to a speech sound (i.e. core stuttering characteristics) as well as syllables in a repeated multi-syllabic word, a repeated phrase or phrase revision, a word fragment, or interjection (e.g. “um”, “err”).

We also measured how fluency was affected over the course of the intervention by including an additional assessment of the change in disfluency immediately after the intervention session on each of the 5 days of intervention.

One researcher completed all disfluency counts. Inter-rater reliability was measured by comparing all speech samples from two participants, selected at random, with counts independently completed by a second researcher. A strong intra-class correlation was found for the inter-rater measurements (ICC =.94, p <.001), indicating a high level of reliability.

Secondary outcome measures included the SSI-4, which provides a standardised and norm-referenced index of disfluency, and the Overall Assessment of the Speakers Experience of Stuttering (OASES: Yaruss *et al*., 2006), a self-assessment tool, which measures the psycho-social impact of stuttering. The latter was used at baseline and at the 6-week post-intervention time point, to avoid violating retest reliability.

Speech naturalness was monitored across the trial as a reduction would be considered a possible adverse effect of stuttering intervention (Martin *et al*., 1984; Onslow *et al*., 1992; Teshima *et al*., 2010). tDCS has been associated with mild and transient adverse effects. Therefore, we also monitored adverse effects related to receiving tDCS such as an itching or tingling sensation at the electrode sites.

### Statistical analysis

It was not possible to perform a power calculation based on previous trials of tDCS in developmental stuttering, as no studies prior to this one were published. Previous intervention studies using tDCS in patients with aphasia reported group differences of medium effect size (e.g. using a sample size of n = 10, Baker *et al*., 2010).

Data were analysed according to the intention-to-treat principle. The distribution of the measurement of speech disfluency at baseline significantly deviated from normal, as is commonly seen in people who stutter (Jones *et al*., 2006). To avoid the need for transformation (which is problematic for reporting confidence intervals in interpretable units, Bland *et al*., 1996), the trial outcomes were defined in terms of change from baseline, which was normally distributed.

The effect of tDCS on the primary outcome measure (change in %ds from baseline) was assessed using a mixed-model analysis of variance (ANOVA), with a between-subjects factor of group (tDCS, sham) and two within-subjects factors of time post-intervention (1 week, 6 weeks) and speech task (reading, conversation). Further ANOVAs for the two groups separately were used to explore significant interactions. The effect of tDCS on the secondary outcome measure of (change from baseline in the SSI-4 score) was also assessed using a mixed-model ANOVA with the between-subject factor of group (tDCS, sham) and a within-subjects factor of time post-intervention (1 week, 6 weeks). As the other secondary outcome measure (change from baseline in OASES) was only acquired 6 weeks post intervention, the effect of tDCS on this measure was assessed using an independent samples t-test between the two groups. For the additional analysis of the effects of tDCS during the 5-day intervention on speech fluency, we entered the change from baseline %ds measured post-intervention on each day into a mixed-model ANOVA. Group (tDCS, sham) was the between-subjects factor and task (reading, conversation) and intervention day (1 to 5) were within-subjects factors.

The means of changes from baseline in %ds, with 95% confidence intervals, were calculated for the tDCS and sham groups separately, along with the differences in these means between the two groups. Cohen’s d was calculated for the effect sizes of the group differences. The change from baseline in %ds was also calculated as a percentage of the median %ds at baseline to estimate the size of the change relative to the baseline rate of disfluency.

## Results

Between October 2014 and February 2016, 71 adult men who stutter were assessed for eligibility for the study. Thirty-four were ineligible either because their stuttering severity was assessed as mild (n = 28), which was below our cutoff of moderate severity, or because they had an additional language disorder (n = 2), or contra-indications to brain stimulation (n = 4). Seven declined to participate. Thus, 30 participants met the eligibility criteria and were recruited. All participants completed the intervention and post-intervention sessions, and were included in all the analyses. The one-week post-intervention session was carried out on average 8 days after intervention and the six-week session at 40 days after intervention. Table 1 shows baseline characteristics, which were well-matched between the tDCS and sham groups.

**Table 1:**
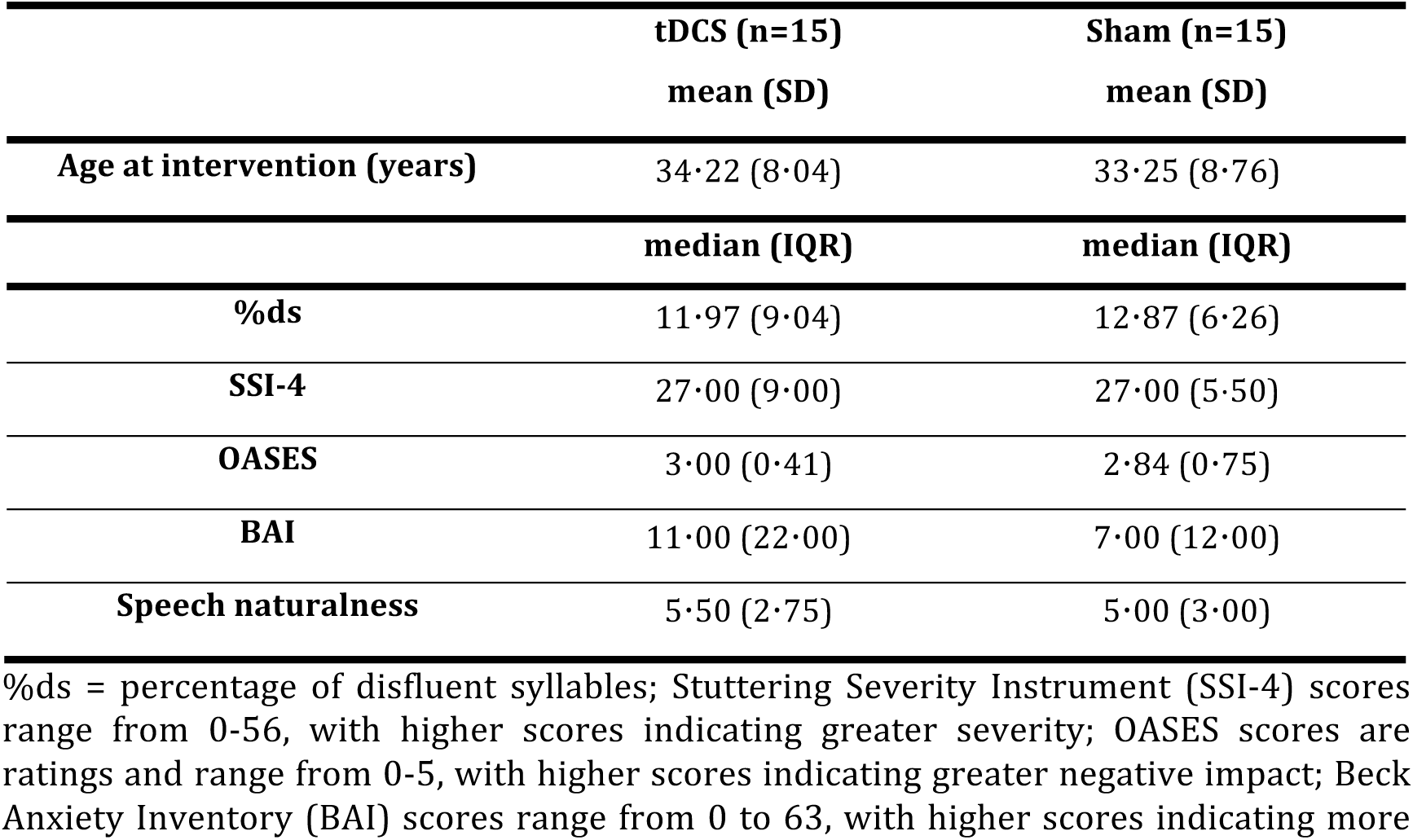
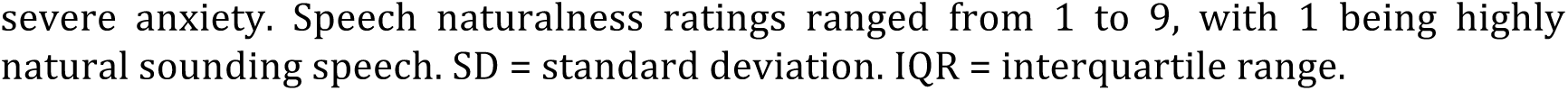
Baseline characteristics.

|  | **tDCS (n=15)** | **Sham (n=15)** |
| --- | --- | --- |
|  | **mean (SD)** | **mean (SD)** |
| **Age at intervention (years)** | 34.22 (8.04) | 33.25 (8.76) |
|  | **median (IQR)** | **median (IQR)** |
| **%ds** | 11.97 (9.04) | 12.87 (6.26) |
| **SSI-4** | 27.00 (9.00) | 27.00 (5.50) |
| **OASES** | 3.00 (0.41) | 2.84 (0.75) |
| **BAI** | 11.00 (22.00) | 7.00 (12.00) |
| **Speech naturalness** | 5.50 (2.75) | 5.00 (3.00) |

%ds = percentage of disfluent syllables; Stuttering Severity Instrument (SSI-4) scores range from 0-56, with higher scores indicating greater severity; OASES scores are ratings and range from 0-5, with higher scores indicating greater negative impact; Beck Anxiety Inventory (BAI) scores range from 0 to 63, with higher scores indicating more
severe anxiety. Speech naturalness ratings ranged from 1 to 9, with 1 being highly natural sounding speech. SD = standard deviation. IQR = interquartile range.

One participant in the tDCS group was an extreme statistical outlier (>3 standard deviations from the group mean) with regard to baseline stuttering, but this participant’s change from baseline scores were not outliers. Data from all participants were included in the primary analysis, according to the intention-to-treat principle. However, we also completed sensitivity analyses (Thabane *et al*., 2013) by re-running all analyses excluding the participant with outlying baseline scores, to evaluate the robustness of the treatment effect. The sensitivity analyses resulted in minimal change to the tDCS group mean, and did not alter the pattern of results regarding the effects of tDCS.

Table 2 shows mean change and confidence intervals per group for all outcomes measures. Figure 2 shows mean change in %ds, the primary outcome measure, for both groups, at both post-intervention time points. Figure 3 shows the changes from baseline disfluency for the two speaking tasks in each group separately.

**Table 2:**
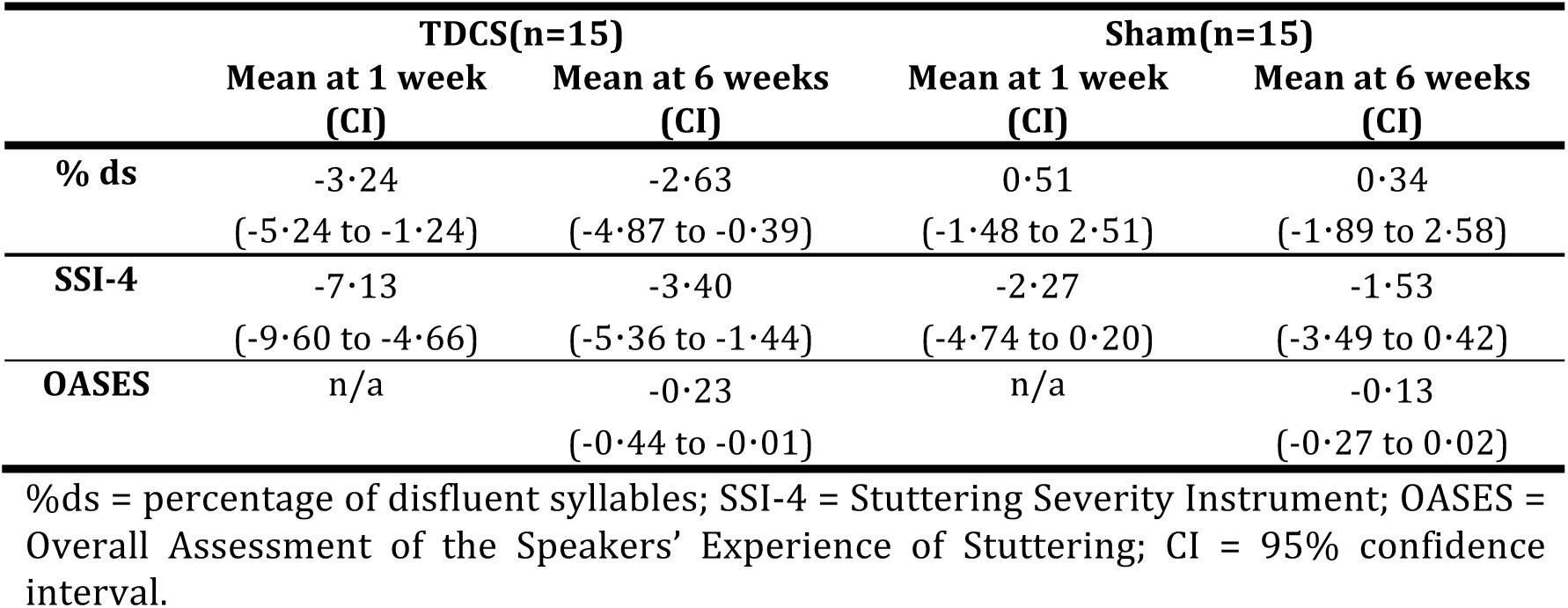
Summary of mean changes from baseline per group for the primary and secondary outcomes

|  | TDCS(n=15) |  | Sham(n=15) |  |
| --- | --- | --- | --- | --- |
|  | Mean at 1 week<br>(CI) | Mean at 6 weeks<br>(CI) | Mean at 1 week<br>(CI) | Mean at 6 weeks<br>(CI) |
| **% ds** | -3.24<br>(-5.24 to -1.24) | -2.63<br>(-4.87 to -0.39) | 0.51<br>(-1.48 to 2.51) | 0.34<br>(-1.89 to 2.58) |
| **SSI-4** | -7.13<br>(-9.60 to -4.66) | -3.40<br>(-5.36 to -1.44) | -2.27<br>(-4.74 to 0.20) | -1.53<br>(-3.49 to 0.42) |
| **OASES** | n/a | -0.23<br>(-0.44 to -0.01) | n/a | -0.13<br>(-0.27 to 0.02) |

%ds = percentage of disfluent syllables; SSI-4 = Stuttering Severity Instrument; OASES = Overall Assessment of the Speakers' Experience of Stuttering; CI = 95% confidence interval.

**Figure 2:**
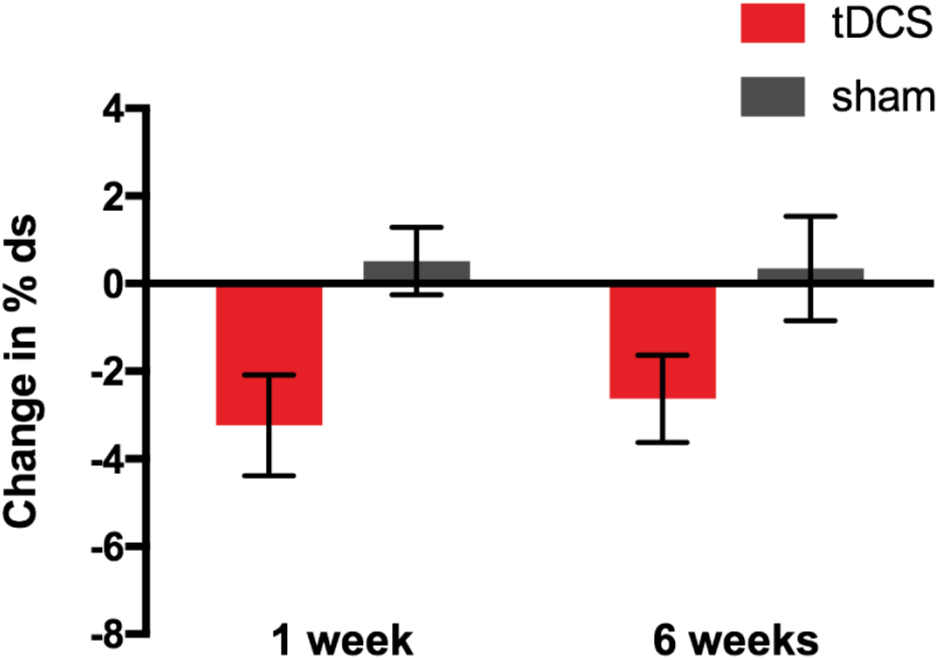
Effect of tDCS on the primary outcome measure: change from baseline in speech disfluency. Bars indicate mean change from baseline in %ds measured at 1-and 6-weeks post intervention averaged across speech samples obtained during reading and conversation. Red – tDCS group; grey – Sham group. Error bars indicate standard error of the mean.

**Figure 3:**
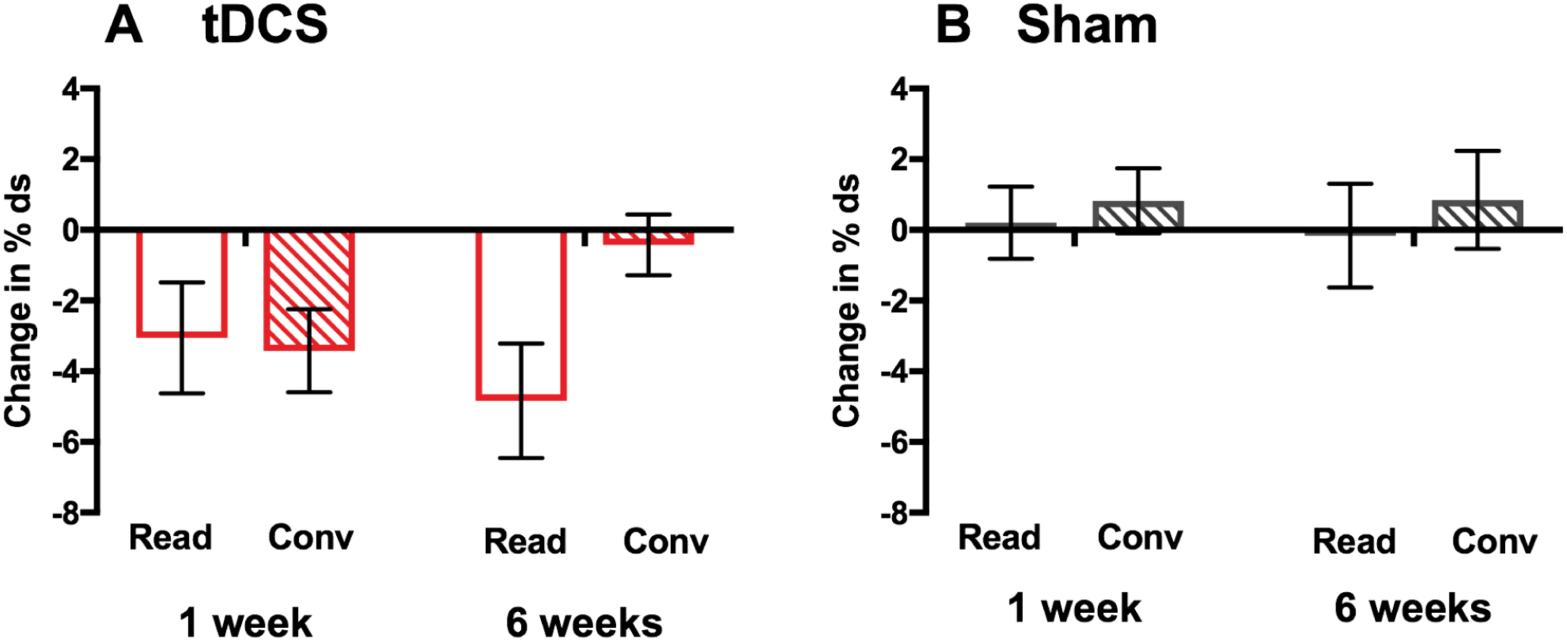
Effect of tDCS on reading and conversation tasks separately. Bars indicate mean change from baseline in %ds measured at 1- and 6-weeks post intervention for the two speaking tasks in the A. tDCS and B. sham groups. Unfilled bars – reading task (Read), striped bars – conversation task (Conv). Error bars indicate standard error of the mean.

For our primary outcome measure, change from baseline in %ds, we found significantly greater reduction in disfluency in the tDCS group relative to the sham group who showed minimal change from baseline (main effect of group, F (1,28) = 7⋅21, p = ⋅012, Cohen’s d = 0⋅98; figure 2). The size of the change from baseline in %ds for the tDCS group was -3.24 at 1 week and -2.63 at 6 weeks after intervention. This change expressed as percentage of the baseline %ds for the tDCS group (11.97%), represents a 27% reduction in disfluency at 1 week and 22% at 6 weeks. In contrast, the change from baseline in %ds for the sham group represented a 4% increase in disfluency at 1 week and a 3% increase at 6 weeks (percentage of their baseline of 12.87%).

Across the two groups, the change in %ds did not significantly differ between the two post-intervention time points (1 and 6 weeks) or between the two speech tasks (reading and conversation) (main effects of task: p = ⋅144; and time-point: p = ⋅774). However, there were significant interactions between task and time-point (F(1,28) = 6⋅62, p = ⋅016) and among task, time-point and group (F(1,28) = 4⋅77, p = ⋅037). The three-way interaction was examined using repeated-measures ANOVA for each group separately with factors of task and time-point. For the tDCS group, there was a significant interaction between task and time-point (F (1,14) = 11⋅13, p = ⋅005; figure 3A) and this was not significant for the sham group (F < 1, p = ⋅786; figure 3B). Examination of the means in Figure 3A suggests that the task by time-point interaction in the tDCS group is due to maintenance of the reduced disfluency relative to baseline for the reading task at 6 weeks (no significant difference between time-points) but a return to baseline disfluency levels for the conversation task (main effect of time-point, p = ⋅034).

For our secondary outcome measure of stuttering severity, we found a significantly greater reduction in SSI-4 score in the tDCS relative to the sham group (main effect of group, F(1,28) = 6⋅31, p = ⋅018; Cohen’s d = 0⋅92; fig. 4A). The reduction in SSI-4 was significantly larger at 1 week compared with 6 weeks post-intervention, for both groups (significant main effect of time point, F(1,28) = 8⋅73, p = ⋅006; fig. 4A). The interaction between group and time point was not significant (F(1,28) = 3⋅94, p = ⋅057).

**Figure 4:**
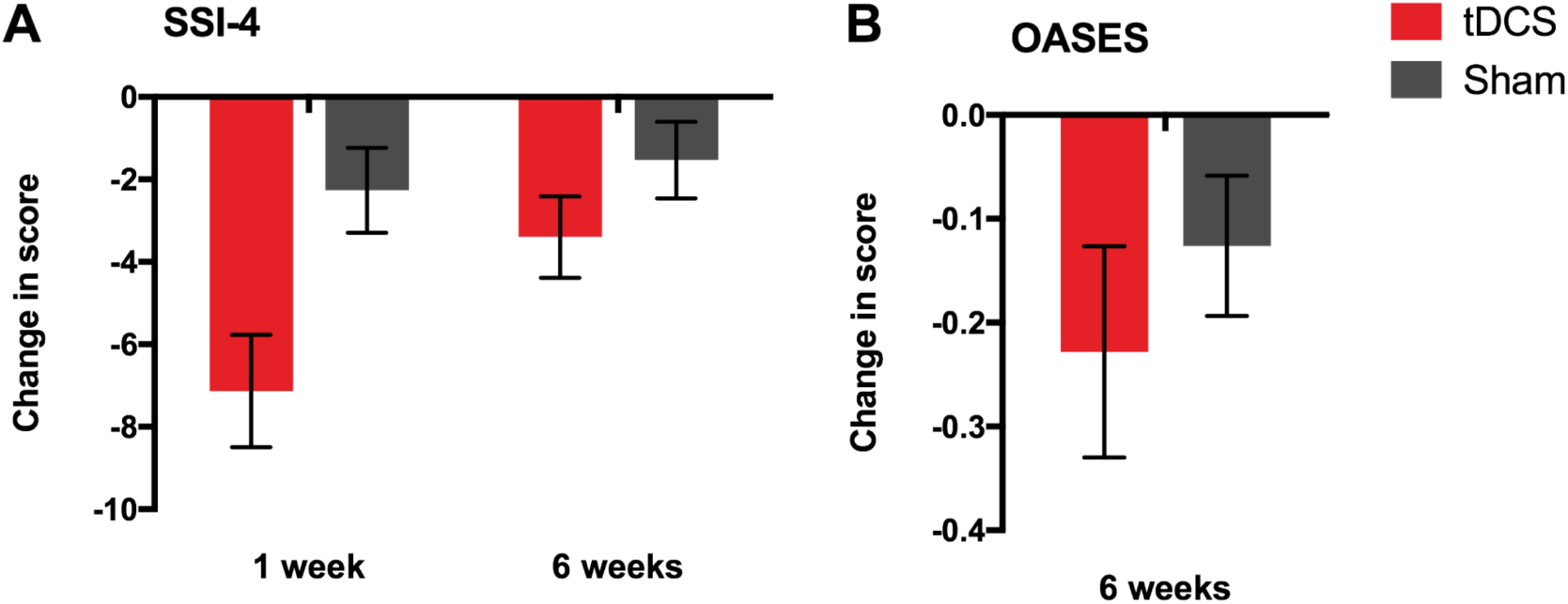
Effects of tDCS on secondary outcomes: change from baseline in SSI and OASES scores. Bars indicate mean change from baseline in A. SSI-4 scores at 1- and 6-weeks post intervention, and B. OASES scores at 6-weeks post intervention, for the tDCS (red) and sham (grey) groups. Error bars indicate standard error of the mean.

For our other secondary outcome measure, change from baseline in the OASES, which was measured only at 6 weeks post-intervention, we found no significant effect of tDCS (independent samples t-test, t (28) = -0⋅84, p = ⋅410; Cohen’s d = 0⋅31; fig. 4B). Examination of the means shown in Figure 4B reveals that the OASES scores were reduced relative to baseline in both groups.

In separate exploratory analyses, we examined the effects of tDCS on change from baseline in %ds during the five-day intervention. There was a significantly larger reduction in %ds over the 5 days of the intervention in the tDCS relative to the sham group (main effect of group, F (1,28) = 9⋅53, p = ⋅005, Cohen’s d = 1⋅13; fig. 5). This main effect did not interact with speech task but in both groups the change from baseline in %ds was significantly greater for the reading compared with the conversation task (main effect of task, F (1,28) = 5⋅36, p = ⋅028). Although the interaction was not significant, it is worth noting that the difference between reading and conversation is clearly evident in the tDCS group and minimal in the sham group (Fig. 5). There was no main effect of time (i.e. day of intervention) nor interaction involving time, task or group.

**Figure 5:**
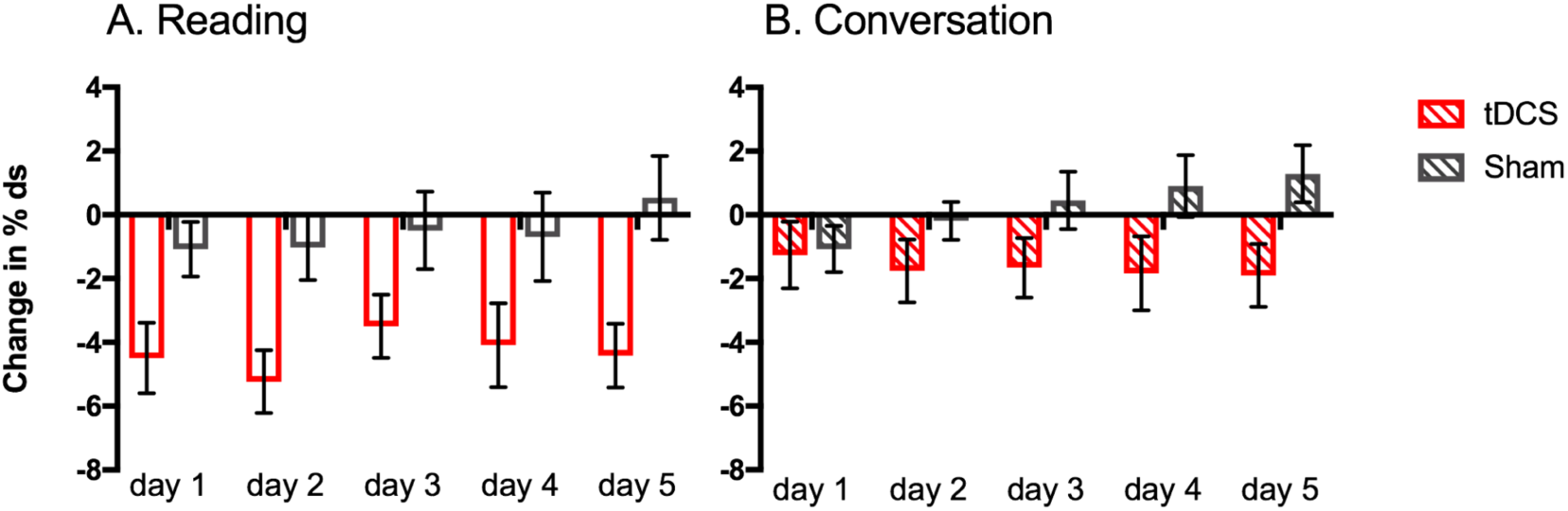
Effects of tDCS on speech disfluency during the 5-day intervention. Bars indicate mean change in %ds from baseline in speech sample during A. reading (unfilled bars) and B. conversation (striped bars) tasks on days 1 to 5 during the intervention for the tDCS (red) and sham (grey) groups. Error bars indicate standard error of the mean.

There were no serious adverse effects during the trial. tDCS adverse effects were limited to the mild symptoms commonly reported in previous studies (e.g. itching and tingling under the electrodes), and did not significantly differ in intensity or frequency of reporting between the tDCS and sham groups. Neither tDCS nor the behavioural intervention alone (the sham condition) affected speech naturalness, when tested immediately after each intervention session, or at the post-intervention time points.

## Discussion

This first randomised controlled trial using tDCS to treat developmental stuttering showed that tDCS in combination with a behavioural fluency intervention significantly enhanced speech fluency compared with sham stimulation. Furthermore, this benefit remained evident at least 6 weeks post intervention. For the primary outcome measure, the percentage of disfluent syllables averaged across reading and conversation tasks at 1 and 6 weeks post intervention was significantly reduced in the tDCS group relative to the sham group. Similarly, the combination of tDCS and fluency intervention significantly reduced scores on a standardised measure of stuttering severity, SSI-4, relative to sham stimulation. This effect also persisted for 6 weeks post intervention. The magnitude and the persistence of improvements for the tDCS group in these outcomes indicate the clinical potential for tDCS as an adjunctive therapy.

The SSI-4 was included in the trial as a widely recognized standardized clinical measure, which provided complementary information to the primary outcome measure regarding fluency disruptions. Specifically, % disfluent syllables is a highly sensitive measure of stuttering frequency, whereas the SSI-4 sacrifices some sensitivity (by conversion to scaled scores), but incorporates important information regarding duration of stuttered moments, and of concomitant features such as tic-like facial or body movements. The effects of tDCS measured by SSI-4 were consistent with those reported for the primary outcome: this composite measure of stuttering symptoms was significantly reduced relative to sham across both post-intervention time points. The sub-scores of the SSI-4 were not analysed separately. However, the size of reductions was larger in the tDCS group than the sham group for all sub-scores (frequency, physical concomitants and duration), except for the duration sub-score at 6 weeks post-intervention, when the groups did not differ.

No significant benefit of tDCS was found for the OASES self-assessment, our other secondary outcome measure. However, both groups showed some reduction in the negative psycho-social impact of stuttering following intervention. We included the OASES as a measure of psycho-social impact, however the assessment has a broader scope, encompassing all domains of health and disability within the World Health Organisation ICF framework (http://www.who.int/classifications/icf/en/). The sub-scores of the OASES were not separately analysed but small reductions were seen for both groups across all sub-scores (assessing general understanding of stuttering, reactions to stuttering, communication in daily situations and quality of life).

Our primary outcome measured change in fluency across the two speaking tasks, reading and conversation, but we were also interested in potential differences in sensitivity to tDCS between the two tasks. In the tDCS group, the significant reduction in disfluency observed 1 week post-intervention was maintained for the reading task at 6 weeks post-intervention but had decreased significantly for the conversation task (i.e. it had returned towards baseline levels). Changes in disfluency for reading and conversation were also considered separately in our additional exploratory analysis of the time-course of tDCS effects during the intervention. We found that tDCS reduced disfluency significantly across the five days of the intervention and that the disfluency decreases were greater for the reading than conversation tasks. It appears therefore that speech samples taken during reading tasks provide a more sensitive measure of disfluency. This may be because it is impossible to avoid difficult words or phrases (i.e. those on which stuttering is predicted) when reading text, whereas during conversation people who stutter commonly report using such avoidance strategies (Riley *et al*., 2004). Nevertheless, fluency during conversation might be considered a more ecologically valid outcome measure of a trial aimed at improving speech fluency. Testing combined tDCS and behavioural therapy paradigms to induce more robust increases in fluency during conversation, for example using a longer intervention period, will be important in the ongoing development of this approach for clinical application.

We chose to use a combination of behavioural interventions that have been shown to immediately, and relatively effortlessly, induce speech fluency in people who stutter. However, these interventions alone are temporary in effect. We predicted that the behavioural interventions would engage brain networks supporting fluent speech, and that tDCS would facilitate the function of these networks during speaking resulting in plastic change and prolonged improvements in fluency. Fluency techniques typically used in speech and language therapy, e.g. ‘fluency shaping’ techniques, involve explicit acquisition of new speaking patterns (Boberg and Kully, 1994; O’Brian *et al*., 2003). tDCS could also be used to modulate learning of these new speaking patterns and either increase the rate of acquisition or prolong the effects of such therapy. Given the promising improvement we found based on these temporary fluency-enhancing techniques, further research is needed to explore the interactions between tDCS and other behavioural interventions, and determine the combinations optimal for clinical benefit.

A side-effect of explicitly learning new speech patterns in fluency therapy, however, can be a reduction in speech naturalness (Metz *et al*., 1990; O’Brian *et al*., 2003; Tasko *et al*., 2007), particularly in the early stages. Reduced naturalness following therapy can result in a more negative listener response than to stuttering itself (Stuart *et al*., 2004), and reduce the maintenance of therapy gains (Onslow *et al*., 1992). The current study, aimed to induce fluent speech immediately and with minimal effort, importantly, not to negatively impact speech naturalness. The maintenance of natural-sounding speech following the combination of tDCS with temporary behavioural fluency enhancement in this paradigm is noteworthy. tDCS as an adjunctive therapy for stuttering would have particular impact if maintenance, or even improvement, of speech naturalness is shown to be a replicable outcome.

One limitation of our study was the application of restrictive eligibility criteria for selection of participants. These eligibility criteria were chosen to maximise sensitivity to change. In a previous feasibility study (Chesters *et al*., 2017), we investigated the use of tDCS alongside choral speech in a group of adults with a wide range of stuttering severity. We found that a single 20-minute session of tDCS during fluent choral speech did not produce statistically significant improvements in speech fluency measured immediately and one hour after intervention. There were several aspects of the study design that may have contributed to the null result, including floor effects in measures of disfluency from participants with very mild symptoms and variability between male and female participants in stuttering symptoms (e.g. Silverman *et al*., 1979) and tDCS effects (Krause *et al*., 2014). The multiple-session design of the current study was more likely to yield positive results, as previous studies using multiple session stimulation have shown that unstable effects following the initial intervention session, can increase and stabilize following subsequent daily stimulation sessions (Baker *et al*.; Reis *et al*., 2009). However, we also made a number of changes regarding our study sample to increase sensitivity, including recruiting participants with at least moderate stuttering to avoid a floor effect, and only male, right-handed, participants to increase group homogeneity. For this first randomised controlled trial these restrictive participant criteria were justifiable in order to ascertain that tDCS has some benefit in stuttering intervention. In future work, it will be important to test effects on a more varied sample of adults who stutter. This will help to establish the clinical scope of the approach, and any systematic differences in effect may also help us to better understand the mechanisms involved in tDCS effects and in developmental stuttering.

The positive outcome of this trial has relevance more broadly to the application of tDCS to disorders of the speech and language system, both acquired and developmental. Our results here are consistent with previous work in aphasia (see reviews by Holland *et al*., 2012; Monti *et al*., 2013; Sandars *et al*., 2016). Of particular relevance to the current trial are two studies showing increased speech motor skill following anodal tDCS over left inferior frontal cortex (Marangolo *et al*., 2013; Marangolo *et al*., 2011), in two small samples of patients with acquired apraxia of speech (three and eight patients respectively). Our larger sample of stuttering participants adds support to the claim that applying anodal tDCS over left IFC can increase speech motor rehabilitation outcomes.

There has been very limited research using tDCS in developmental disorders of communication, perhaps due to an understandable caution regarding interacting with neuro-plastic processes during childhood. However, tDCS has an interesting potential for augmenting the limited therapeutic outcomes for adults living with persistent developmental difficulties, the impact of which can be considerable (Clegg *et al*., 2005; Craig *et al*., 2009; Tanner, 2009). One study found that tDCS over area V5/MT combined with a five-day course of reading therapy improved reading speed and fluency in adults with developmental dyslexia, with benefits persisting one week after the intervention (Heth *et al*., 2015). To our knowledge, there are no other studies in adults with developmental disorders of communication, and none in developmental disorders of speech. Our study suggests that tDCS may be usefully applied to persistent developmental communication disorders, and may have particular value where behavioural therapies alone have failed to produce lasting positive outcomes.

In summary, we found that daily application of 20 minutes of 1-mA anodal tDCS over the left IFC combined with tasks performed under choral and metronome-timed speaking conditions for 5 consecutive days improved speech fluency in 15 men who stutter. Another 15 men who stutter showed no change in speech fluency from the same behavioural intervention paired with sham stimulation. These positive findings provide encouragement for future research in developmental stuttering and other disorders of speech and language. Clinical interventions could be extended to use noninvasive brain stimulation in combination with established speech therapy methods including those aimed at reducing the negative impact of living with these conditions. Brain stimulation using tDCS has very moderate costs, and the devices are simple to use, requiring minimal training. Using tDCS stimulation to improve the efficacy of a therapy could reduce the number of sessions required by an individual, offering savings and allowing more individuals to be treated. Furthermore, it could improve outcomes and prevent relapses. Further work is needed, however, to investigate the limitations of this method, its underlying mechanisms and the optimal tDCS parameters for increasing fluency.

## Contributors

JC, RM and KEW designed the study. JC acquired the data. JC and KEW completed the statistical analysis. JC interpreted the data, with input from RM and KEW. JC wrote the first draft of the manuscript, which was edited by all authors. All authors gave final approval of the version to be published.

## Acknowledgments

We would like to thank all of our volunteers for taking part in the study. We also thank Lisa Bruckert, Anthony Chesters, Charlotte Coyte and Kate Heywood for their assistance with data collection and speech data coding, and Faraneh Vargha-Khadem for access to testing facilities. A Medical Research Council Clinical Research Training Fellowship, MR/K023772/1, to JC, and a Medical Research Council Career Development Fellowship, G1000566, to RM, funded this work.

